# Characterization of a novel humanized heavy chain antibody targeting endogenous retroviruses with anti-lymphoma activity

**DOI:** 10.1101/2024.01.17.576027

**Authors:** Filippo Spriano, Luciano Cascione, Jacopo Sgrignani, Nikolai Bendik, Sara Napoli, Giulio Sartori, Eleonora Cannas, Tao Gong, Alberto J. Arribas, Marco Pizzi, Davide Rossi, Davide F. Robbiani, Andrea Cavalli, Francesco Bertoni

**Affiliations:** Institute of Oncology Research, USI, Bellinzona, Switzerland; SIB Swiss Institute of Bioinformatics, Lausanne, Switzerland; Institute for Research in Biomedicine, USI, Bellinzona, Switzerland; Department of Medicine, Cytopathology and Surgical Pathology Unit, University of Padova, Italy; Oncology Institute of Southern Switzerland, Ente Ospedaliero Cantonale, Bellinzona, Switzerland

## Abstract

Lymphomas continue to pose therapeutic challenges, with a considerable portion of patients facing refractory disease. This study focuses on Diffuse Large B-cell Lymphoma (DLBCL), the most prevalent lymphoma type. Within the human genome, transposable elements (TEs), particularly Human Endogenous Retroviruses (HERVs), constitute a significant yet understudied portion. Among HERVs, the HERV-K family, specifically HERV-K113 and HERV-K115, has intact open reading frames. Epigenetic regulation tightly controls HERV expression, and aberrant expression has been observed in various cancers, including lymphomas.

This research investigates the potential of HERV-K as a therapeutic target in DLBCL. The study encompasses comprehensive methods, including RNA extraction, PCR detection, flow cytometry, immunoblotting, peptide prediction, phage display, surface plasmon resonance, ELISA, antibody-dependent cell-mediated cytotoxicity, internalization assays, and bioinformatic analysis. Results reveal the presence and expression of HERVs in lymphoma patients and cell lines, with the HERV-K envelope protein identified as a crucial contributor to lymphoma cell growth. Moreover, the study identifies immunogenic regions of HERV-K, leading to the development of a humanized camelid nanobody (FF-01) with potential therapeutic applications. Furthermore, bioinformatic analysis differentiates DLBCL subgroups based on TE expression, providing insights into prognostic variations. Patients with high HERV-K113 expression show activation of pathways related to antiviral responses, suggesting a viral mimicry state.

In conclusion, the study highlights the clinical relevance of HERVs in lymphomas, proposing them as novel therapeutic targets. The newly developed nanobody FF-01 demonstrates anti-lymphoma activity through antibody-dependent cellular cytotoxicity and internalization. This research opens avenues for exploring endogenous retroviruses as targets for immunotherapy in lymphomas, showcasing the potential of FF-01 as a promising candidate for further investigation.

## Background

Lymphomas are among the ten most frequent tumors in adults and the top three in children and young adults, and despite the significant improvements in their management, still too many individuals die of their diseases ^1-3^, indicating the need for novel therapeutic agents. Diffuse large B-cell lymphoma (DLBCL) is the most common type of lymphoma, accounting for approximately 30% of all cases in adults and 60% in the younger population ^2,4^. Although more than half of DLBCL patients can be cured, about 40% of patients show refractory disease or relapse after an initial response. The outcome remains poor for many of these cases, even after introducing the most recent antibody-based or cellular therapies ^5^.

The human genome is composted for the 45% of transposable elements (TEs) that can be divided into four types: long interspersed elements (LINEs), short-interspersed elements (SINEs), Human endogenous retroviruses (HERVs) and DNA transposons ^6^. Among these, HERVs derive from exogenous retrovirus infections incorporated in germline DNA millions of years ago ^7,8^. Although they encompass a considerable fraction of our genome (up to 8%) ^6,8^, their biological role is understudied. Indeed, HERVs are discarded mainly by the default transcriptome data mining pipelines since their DNA sequences primarily appear multimapped or ambiguous ^8,9^.

HERV-K is a human-specific HERV family with intact open reading frames (ORF) ^7,10^. Phylogenetically, it belongs to the betaretrovirus-like supergroup as its sequence is most similar to mouse mammary tumor virus (MMTV)^11^. HERV-K family comprises ten subgroups representing separate germline infections, termed human endogenous MMTV-like (HML)-1 to HML-10 (human endogenous MMTV-like). HERV-K (HML-2) group is the most recent retrovirus to colonize the human germ line ^12^. Members of the HML-2 subgroup possess complete open reading frames (ORFs) with coding capability and are the most studied concerning diseases ^7,10^. Two HERV-K elements, part of the HML-2 subgroup, HERV-K113 and HERV-K115, are fascinating since they represent full-length proviruses. HERV-K113 and HERV-K115 have complete open reading frames for all viral proteins ^13^. HERV-K113 and HERV-K115 can be found in the genome as single copies located on chromosome 19p13.11 and 8p23.1, respectively ^14^. Their frequencies heavily vary based on a population base, with high prevalence in Africans (up to 30-40%) and lower in Caucasians and Asians (up to 10%) ^14,15^.

Epigenetic mechanisms tightly regulate HERV expression, and usually, in normal tissues, HERVs are heavily methylated and not expressed ^8,16^. In contrast, physiological expression of HERV elements has been observed in embryonic stem cells ^8,17^. Indeed, HERVs can contribute to normal cellular and tissue function, as demonstrated by the Syncytin-1 and Syncytin-2 envelope proteins, encoded by the HERV-W and HERV-FRD, respectively ^18,19^. These proteins play a crucial role in facilitating the fusion of trophoblasts, mediated by their expression on the surfaces of both cytotrophoblasts and syncytiotrophoblasts ^8,18,19^.

An aberrant expression of several HERVs has been demonstrated both in human cancer cell lines and primary tumor tissues, such as breast cancer, renal cell carcinoma, ovarian cancer, melanoma, and myeloproliferative disorders, in which high levels of HERV-K RNA and proteins have been described ^8,20,21^. HERV-K proteins are believed to contribute to tumor cell growth and survival, but little is known about the underlying mechanisms. HERV element expression has recently been observed in lymphoma and leukemia cell lines and patients. ^22-24^. Moreover, elements of the HERV-K family have been identified in the plasma of lymphoma patients ^25^. Here, we explored HERV-K as a therapeutic target in B-cell lymphomas.

## Methods

### Cell lines

Cell lines were cultured according to the recommended conditions, as previously reported ^26,27^. Media were supplemented with fetal bovine serum (FBS, 10%) plus 0.1% Penicillin-Streptomycin 100X (Euroclone, ECB3001D). Short tandem repeat (STR) DNA fingerprinting confirmed the cell line’s identity using the Promega GenePrint 10 System kit (B9510)^28^. Cells were tested to verify mycoplasma negativity using the MycoAlert Mycoplasma Detection Kit (Lonza, Visp, Switzerland).

### RNA extraction and envelope quantification

Total RNA was extracted with TRIzol Reagent (Thermo Fisher Scientific, Waltham, Massachusetts, US). DNAse treatment was performed (Sigma-Aldrich, St. Louis, Missouri, US). Reverse transcription and quantitative real-time PCR were performed with the QuantiFast SYBR Green PCR Kit (Qiagen, Hilden, Germany). Since the endogenous retrovirus envelope comprises only one exon, negative control without retro transcription was applied to infare the DNA contamination.

Primers used for HERV env q-PCR are described by Gröger et al. ^29^. GAPDH was used as a negative control.

### HERVK-113 PCR detection

Genomic DNA was extracted from cell lines using the DNeasy Blood & Tissue Kits (Qiagen, Hilden, Germany) and amplified using the strategy described by Moyes et al. ^30^.

### Flow cytometry

Approximately 500’000 cells for each condition were harvested, washed, and incubated with primary antibody 30’ in ice. Similarly, cells were incubated with a secondary antibody for 30’ in ice. Flow cytometry was performed with BD FACSCanto II (BD Biosciences, California). Analysis was performed with FlowJo software. Anti-HERV-env Ab (MBS9216561, Mybiosource); Isotype control Ab (ab37415, Abcam).

FACS buffer: PBS, 1% BSA, 0.1% sodium azide

### Immunoblotting

Protein extraction, separation, and immunoblotting were performed as previously described ^31^. The following antibodies were used: anti-HERVK env (MBS9216561, MyBioSource), anti-GAPDH (FF26A) from eBioscience, secondary mouse (NA931V) and rabbit (NA934V) antibodies from GE healthcare. Data were analyzed with Fusion Solo software (Vilberg, France).

### Peptide prediction

The HERV-K113 sequence and the prediction about the localization of protein domains and putative glycosylation points were retrieved from the Uniprot database ^32^ using Q902F9 as protein entry. The AlphaFold model was obtained from the AlphaFold Protein Structure Database ^33,34^. Custom peptides were purchased from GeneScript.

### Phage display

The recombinant antibodies utilized in this investigation were sourced from the Geneva Antibody Facility, accessible at https://www.unige.ch/medecine/antibodies/. In summary, a VHH antibody phage library underwent selection against biotinylated target peptides immobilized on Streptavidin-coated magnetic beads. The phages bound to the target were subsequently amplified, and this selection process was iterated for an additional two rounds. Following the conclusive selection round, the chosen antibody phages were transformed into recombinant VHH-Fc antibodies and expressed as cell supernatant in HEK293 cells.

### Surface plasmon resonance

Surface plasmon resonance (SPR) experiments were run on a Biacore 8000K instrument. Streptavidin was immobilized on the CM5 surface by a standard amino coupling. Streptavidin was used to immobilize pept1 and pept2 by the biotin moiety bounds to the peptide n-terminal ends. Five increasing concentrations of the nanobodies 6.25, 12.5, 25, 50, and 100 nM were injected using a single-cycle kinetics setting. The running buffers were ten mM HEPES pH 7.4, 150 mM NaCl, three mM EDTA, and 0.005% Tween-20. Subsequently, curve fitting was executed using Biacore Insight Evaluation Software (version 5.0.18). The fitting successfully passed the quality control assessment integrated into the software.

### ELISA

Ninety-six well plates (Nunc MaxiSorp Flat-Bottom Plate, 44240421, Invitrogen) were coated with NeutrAvidin (2μg/ml; Thermo Fisher, Massachusetts, US) overnight at room temperature (RT). Plates were washed and subsequently coated with biotinylated peptides for 1 hour at RT. Plates were washed, blocked, and incubated for 1 hour with plasma (1:50 dilution). Peroxidase-conjugated anti-human IgG (Sigma-Aldrich, St. Louis, Missouri, US) was added (1 hour, RT). Finally, substrate (Pierce TMB Substrate Kit, Thermo Fisher Scientific, Massachusetts, US) was administered. Absorbase (450nm) was red with BioTek Cytation 3 (Agilent, California, US).

The cut-off for positivity was calculated with the formula: cut-off = (mean + 3 sd)^35^, where mean and sd were equal to the mean and the standard deviation of the values below the change point. The change point was calculated with R. The analysis was done for each plate, and the median value for each experiment was used.

### Antibody-dependent cell-mediated cytotoxicity

Antibody-dependent cell-mediated cytotoxicity (ADCC) was performed with the ADCC Reporter Bioassay, Core Kit (Promega, Wisconsin, USA), following the manufacturer’s protocol, with an effector-to-target cells ratio of 3 to 1 (45,000 Effector and 15,000 target cells). FF-01 was compared to a non-binding nanobody (FF-00) or an isotype control antibody. ADCC induction was evaluated after 6 or 14 hours of coculture.

### Internalization

Internalization was assessed using the Incucyte Fabfluor-pH Antibody labeling reagent (Sartorius, Germany), following the manufacturer’s protocol. Cells (30,000) were seeded in a 50 μl volume for each well of 96-well plates coated with poly-L-ornithine. Cells were allowed to adhere to the plate bottom for at least one hour and subsequently incubated with 50 μls containing the antibodies (1 μg/ml) and the Fabfluor antibody at a 1:6 molar ratio. Plates were inserted in the live cell imaging Incucyte, and images were taken every 45 minutes. Analysis was performed with the internal Incucyte analysis software as the red area was divided on phase confluence normalized on time zero.

### Envelope downregulation

Transient knockdown was performed using the Amaxa 4D Nucleofector system (Lonza, CH). SiRNAs were designed with the Sfold ^36^ software and the tool provided by Dharmacon. Two different siRNAs were designed and tested, compared to a nontargeting siRNA as control, and BLOCK-iT Fluorescent Oligo was used to check the electroporation efficiency. Protocols according to the SG Cell Line 4D-Nucleofector X Kit L (Lonza) were followed. Briefly, 2 × 10^6^ cells were prepared and resuspended in 100 μl SG solution with 800 pmol siRNA. Flow cytometry confirmed electroporation efficiency 48 hours after nucleofection via the BLOCK-iT control. Cell proliferation was evaluated multiple times (every 24 hours) up to 72 hours. Protein downregulation was determined 72 hours after the beginning of the experiment.

### Bioinformatic analysis

The fastq files for EGAD00001003783 ^37^ and GSE71702 ^38^ datasets were downloaded and separately processed, including quality control, mapping, and gene qualification using Gencode v22. The aligned BAM files were inputted into Telescope (v1.0.3) ^9^. The raw counts’ matrices of the two datasets were merged and filtered: transposable elements with more than five observations within a minimum sample threshold were retained (cardinality of the smallest group). At the locus-specific level, we quantified HERVs expression using an in-house workflow based on Telescope ^9^ and its GTF annotation file (∼15000 HERVs). The obtained raw read count matrix served as input for all downstream analyses presented in this study.

Only features with more than five observations within a minimum sample threshold were retained. Counts were normalized with the TMM algorithm in edgeR ^39^, and batch effects were removed using the removeBatchEffect function of limma ^40^. MDSPlot on the log2 transformed counts per million (lcpm) were obtained using the plotMDS function in limma for visualization. Samples were clustered with the k-means algorithm with k=4 in the stats R package (version 3.6.2). Fisher’s exact test was used to test each cluster against the cell-of-origin (COO). To compare the HERV clusters to the COO-based classification, we generated alluvial plots using ggalluvial (v0.12.3). Survival analysis was done with the survival R package (v3.1-12) and group-level comparisons using the log-rank test. Kaplan-Meier survival plots were created using survminer (v0.4.9) and ggplot2 (v3.3.6). All analyses were performed using Bash, R (v4.2.2), and the Bioconductor package manager (v1.30.19). An in-house script has been developed to quantify the expression of LMP2A and LMP2B genes: we count the reads mapped to the exons of the two genes according to the gtf of the EBV genome.

## Results

### Transposable elements transcripts identify clinically relevant DLBCL subgroups

We first assessed the expression of TEs, including HERVs, in two publicly available RNA-seq datasets (EGAD00001003783 ^37^ and GSE71702 ^38^) of DLBCL and B cells from healthy individuals (centroblast, centrocytes, memory B cells, naïve B cells, TPC). An unsupervised multidimensional scaling (MDS) plot showed that the expression of TE could separate neoplastic from healthy samples (Figure 1A). These data demonstrated that tumoral and normal specimens have distinct TE and HERV expression profiles.

**Figure 1.**
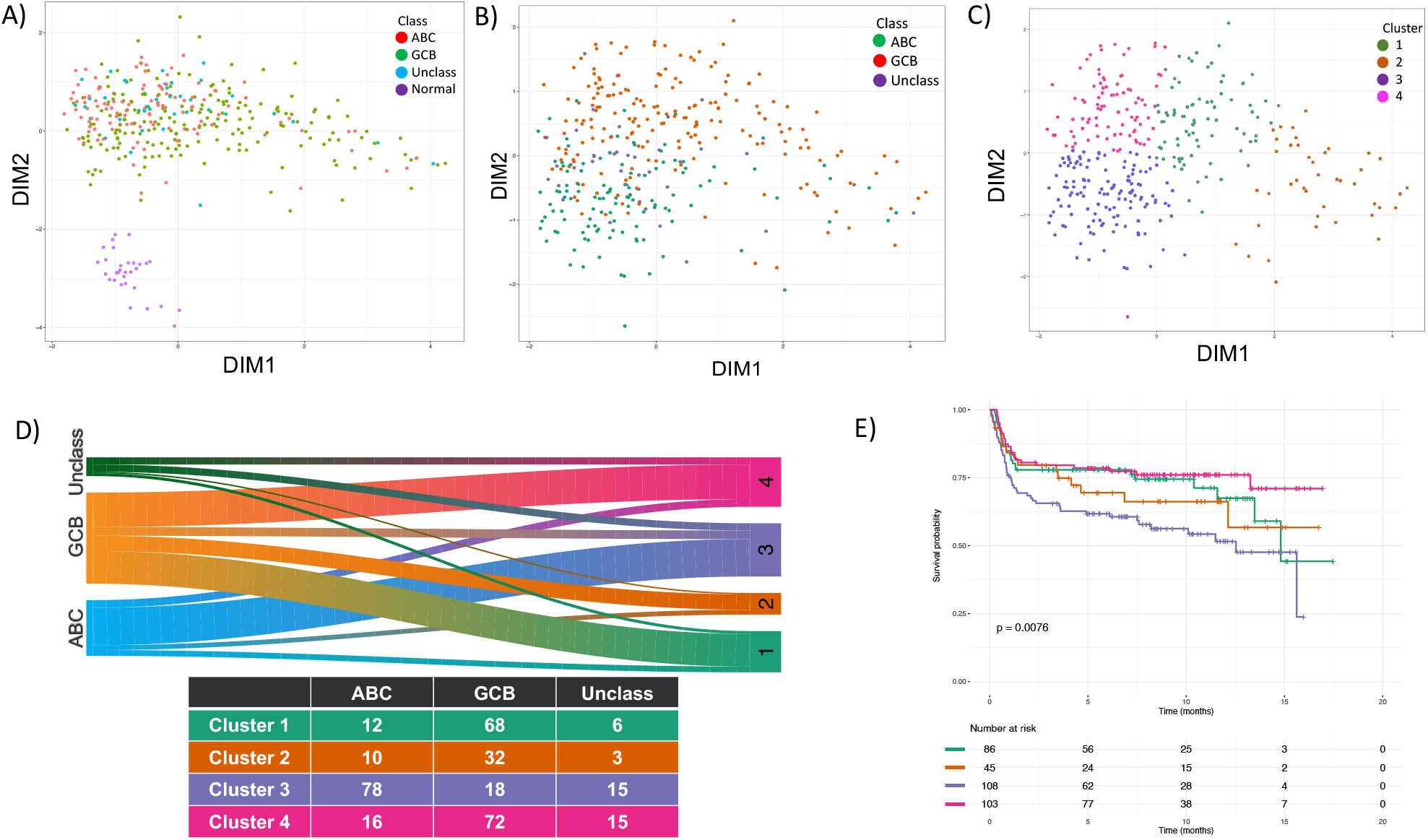
Transposable elements are expressed in DLBCL clinical specimens. (A) Multidimensional scaling (MDS) plot showing normal and tumoral samples clustering differently based on TE expression. (B) Multidimensional scaling (MDS) plot on tumoral samples based on HERV expression. Patients are colored differently based on the ABC, GCB, or unclassified DLBCL subtype. (C) Multidimensional scaling (MDS) plot on tumoral samples clustering them in 4 groups based on HERV expression. (D) Alluvial plot showing the distribution of ABC, GCB, and unclassified patients in the four clusters. (E) The survival curve of the patients is divided into four different clusters

DLBCL is a heterogeneous group, and it contains two main subgroups, the activated B cell-like (ABC) and the germinal center B cell-like (GCB) DLBCL, based on the supposed cell of origin (COO) ^4,41^. When we focused on tumor samples only, the MDS plot separated DLBCL samples based on their COO (Figure 1B). Via k-means, DLBCL samples were subdivided into four clusters (Clusters 1/2/3/4) based on TE expression (Figure 1D). ABC DLBCL samples were mainly present in cluster 3, while GCB DLBCL samples were primarily contained in clusters 1, 2, and 4, with cluster 1 more enriched of double hit/dark zone signature DLBCL (Table S1). The four clusters differed in clinical outcome among patients treated with standard chemo-immunotherapy R-CHOP (rituximab, cyclophosphamide, doxorubicin, vincristine, prednisolone). Cluster 3 identified patients with the worst outcome, followed by Cluster 2 and Cluster 1, while Cluster 4 patients had the best outcome (Figure 1E).

We next focused on the HERV-K (HML-2) family since it can encode an intact ORF. The RNA expression of the HML-2 HERVs family could distinguish normal and tumoral samples (Figure S1). Looking for differences in RNA expression between tumoral and normal specimens, we identified many HERV-K significantly upregulated in tumors, with K113 being one of the most differentially expressed (Figure 2).

**Figure 2.**
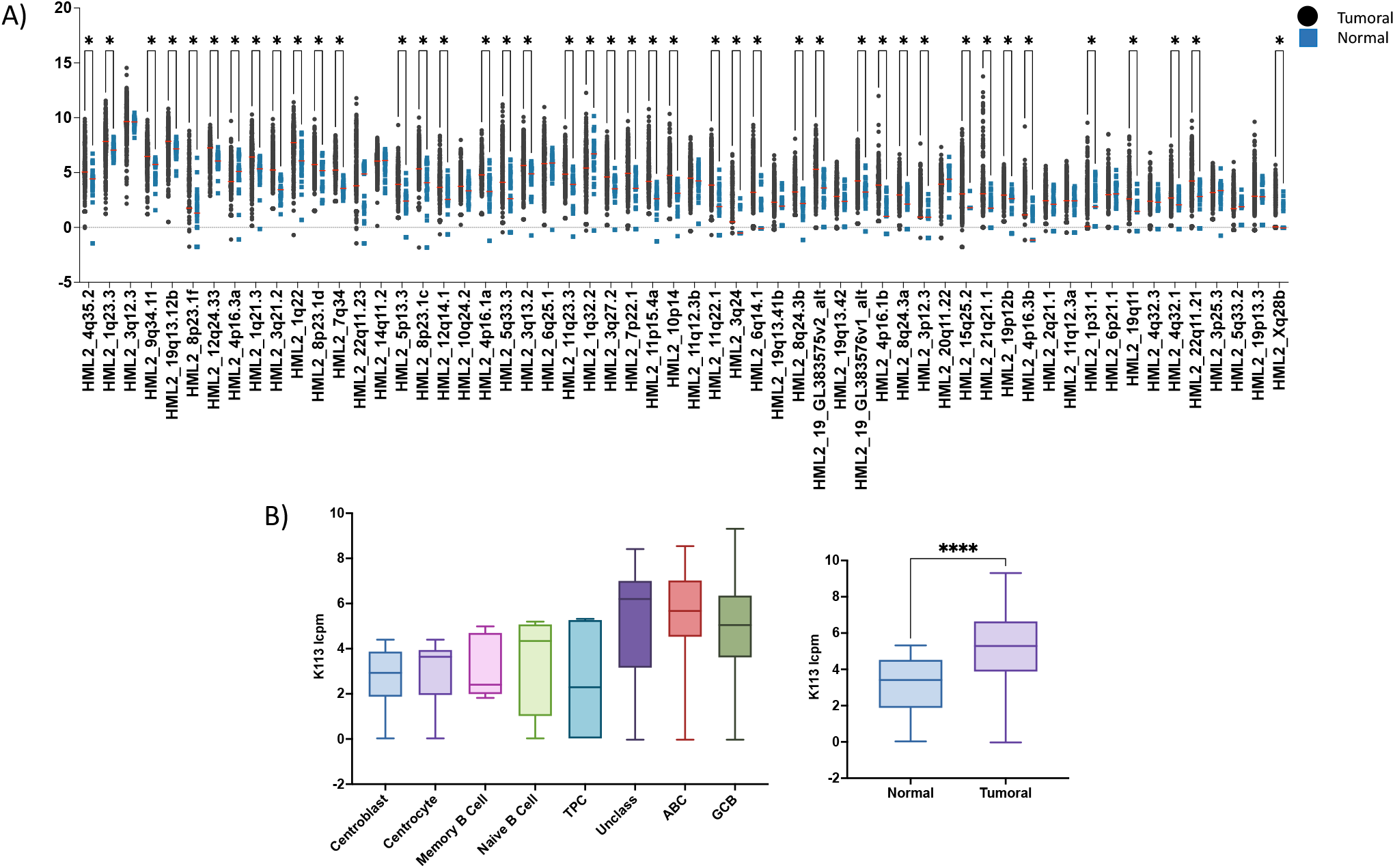
HML-2 expression in patients vs healthy samples. **(A)** HERV-K (HML-2) RNA expression in tumoral versus normal samples. **(B)** HERV-K113 RNA expression in tumoral versus normal samples. * = p value <0.05. lcpm = log counts per million

We divided patients’ samples (EGAD00001003783) between high and low K133 expression. These patients also represent the majority (more than 90%) of patients with an overall high expression of HML-2 HERVs that retain the env gene. We identified that patients with high K113 RNA levels have activation in pathways related to Interferon, toll-like receptors, JAK/STAT, interleukins, and viral infection, suggesting an active state of a phenomenon known as “viral mimicry” in high-expression patients (Figure S2, Table S2). HERV expression in tumoral samples has been linked to a viral mimicry state where cells activate an antiviral response triggered by the expression of endogenous retroviruses ^42^.

Similar results obtained in patients were obtained in a panel of 47 B cell lymphoma cell lines, derived from ABC DLBCL, GCB DLBCL, mantle cell lymphoma (MCL), marginal zone lymphoma (MZL), chronic lymphocytic leukemia (CLL), Burkitt lymphoma (BL) and primary mediastinal B-cell lymphoma (PMBCL). TEs expression identified three clusters of samples. One contained mostly GCB DLBCL, the second had all MCL and six GCB DLBCL, and the third cluster contained MZL, the majority of ABC DLBCL and CLLs (Figure S3). GCB DLBCL was divided into two clusters reminiscent of what was observed in patients. The different clusters did not correlate with gender or EBV status, as assessed via LMP2A and LMP2B expression (Figure S3B/C/D).

### HERV-K envelope protein sustains the growth of lymphoma cell lines

As mentioned above, members of the HERV-K family can present an intact ORF and code for the envelope (env) protein, which can be expressed on the cell surface ^10^. Thus, we studied seventeen DLBCL cell lines by flow cytometry using a monoclonal antibody that recognizes the env of multiple HERV-K family members ^43^. The levels of env cell surface expression varied among the DLBCL cell lines, and they were confirmed via immunoblotting and qPCR (Figure 3A, B; Figure S4). Among the cell lines tested, RCK8 exhibited the highest cell-surface expression of the HERV-K env protein (Figure 3C). HERV-K113 is integrated into the human genome at 19p13.11, and it is possible to detect its presence by PCR ^14,15,30^. RCK8 was the only cell line that showed the genomic insertion (heterozygous) of HERV-K113 (Figure S5). Env expression in the other cell lines was not paired with demonstrated genomic insertion at 19p13.11, and they likely reflected the presence of HML-2/HERV-K family members other than K113, still able to produce an envelope protein.

**Figure 3.**
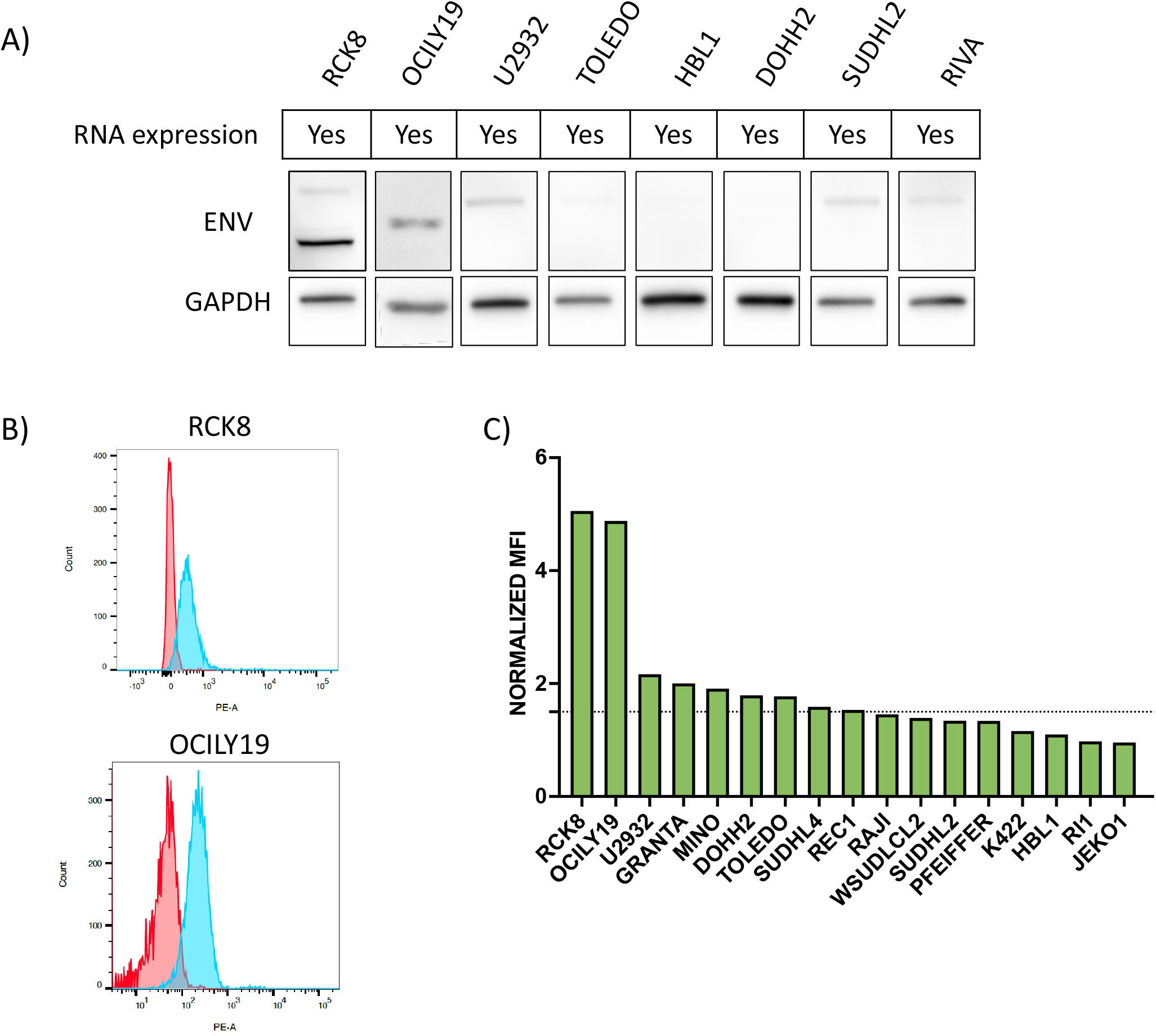
HERV-K envelope protein is expressed in DLBCL cell lines. (A) Immunoblot for HERV-K env expression in DLBCL cell lines with (B) Surface expression of HERVK envelope in lymphoma cell lines, with (C) the individual quantification (MFI=median fluorescence intensity). Anti-HERVK envelope antibody showed envelope surface expression in some lymphoma cell lines. Rabbit polyclonal IgG was used as a negative control.

We could detect different-sized envelope proteins in the other lymphoma cell lines by immunoblotting. This could reflect the expression of additional HERV-K family members or that the envelope protein is differentially post-translational modified. Interestingly, the two cell lines expressing the envelope on the surface showed a lower band size.

To determine whether the expression of HERV-K env is biologically relevant for lymphoma cells, we silenced it with two siRNAs in RCK8 cells (Table S3). Each env-specific siRNA reduced env expression by more than 50% over control (Figure 4A/B) and the reduction was associated with a decrease in cell proliferation rather than a decrease in viability (50% decrease in proliferation at 72 hours after siRNA treatment (Figure 4C, D). These data showed that HERV-K env is expressed on DLBCL cells, and its expression contributes to lymphoma cell growth.

**Figure 4.**
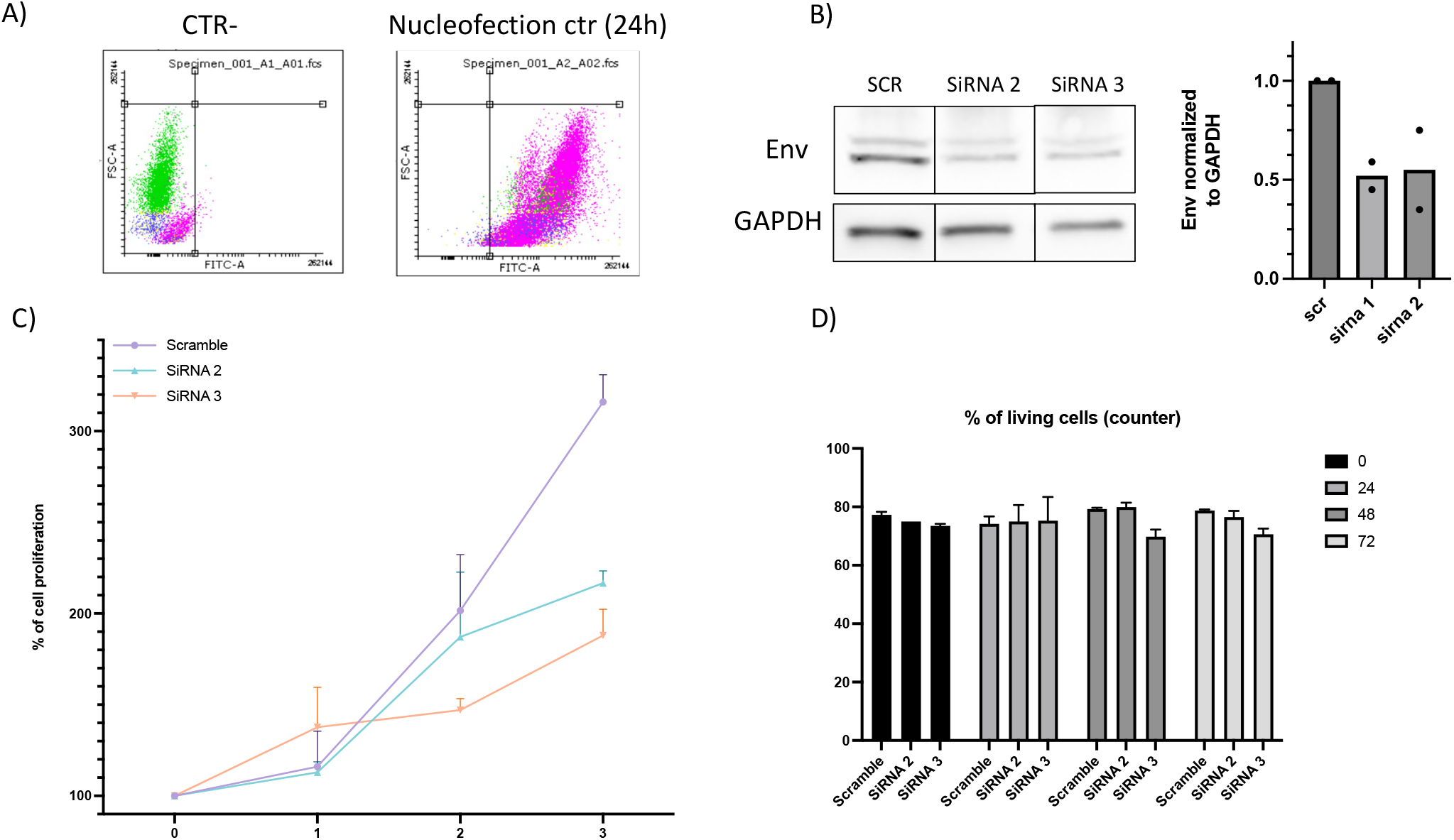
HERV-K envelope protein sustains the growth of lymphoma cell lines. (A) Nucleofection efficiency control shows around 100% of nucleofection efficiency. (B) Downregulation control after siRNAs compared to Ctr treated cells (SCR = scramble siRNA sequence). GAPDH was used as a housekeeping gene. (C) Proliferation curve of cell lines treated with CTR siRNA (scramble) or siRNA2 or siRNA3 (D) Percentage of living cells after siRNA2 or siRNA3 treatment.

### Serum of lymphoma patients contains antibodies targeting HERV-K env

Because env expression is low-to-undetectable in healthy tissues ^44,45^ but upregulated in lymphoma and other cancers ^44,45^, we wondered whether HERV-K could serve as a target for immunotherapy. Since no experimentally determined 3D structure was available in the protein data bank (PDB) ^46^, we predicted a HERV-K113 env model using AlphaFold ^33,34^. In agreement with data reported for similar proteins ^47^, the model (Figure 5A/B) included two main regions, one predicted to be transmembrane or cytosolic and one extracellular on the surface. Examination of the model’s extracellular region identified five linear regions with potential for antibody binding, devoid of putative glycosylation sites and cysteine residues (Figure 5A).

**Figure 5.**
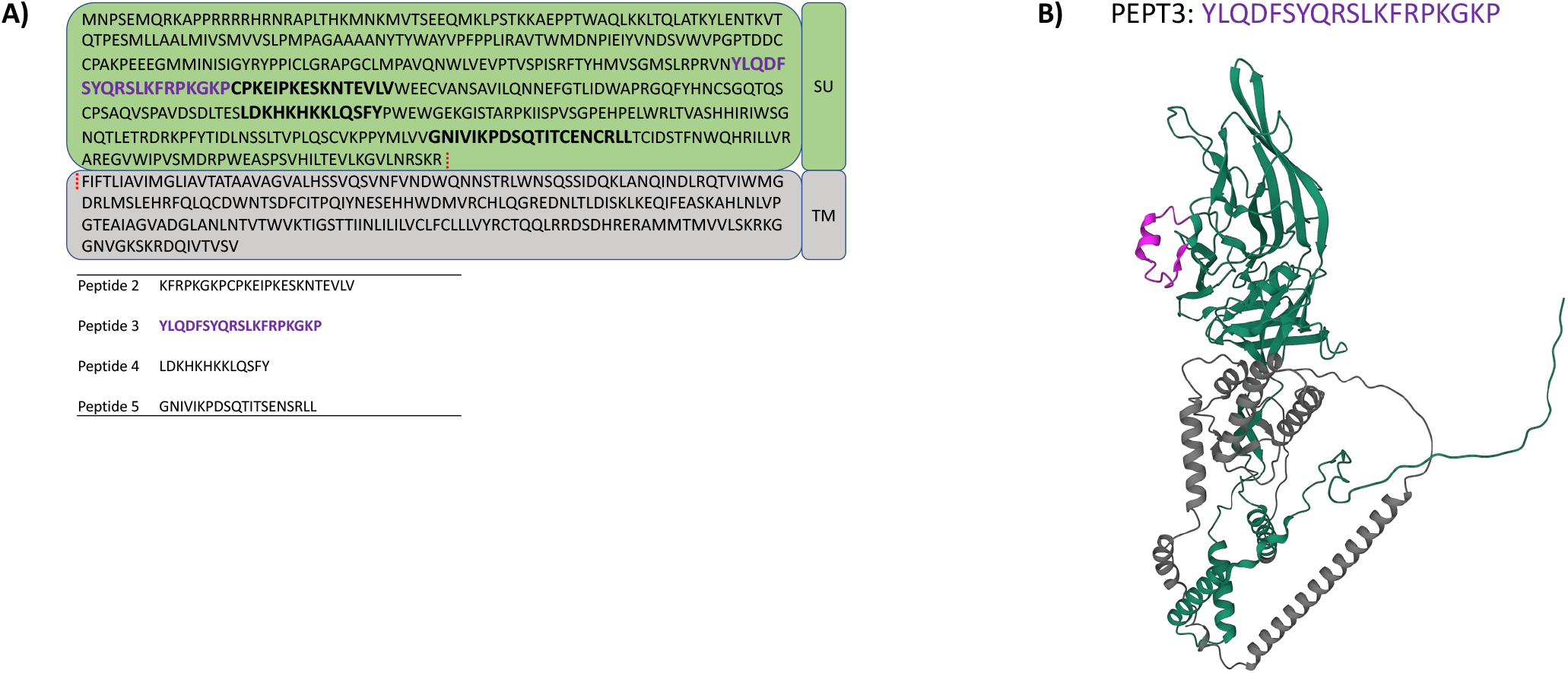
HERV-K envelope surface epitopes. (A) HERV-K113 protein sequence. In Bold, highlight the sequences corresponding to the identified peptides (listed below). SU, surface portion. TM, transmembrane, or cytosolic portion. (B) 3D protein structure modeling of HERV-K113. The peptide three (PEPT3) sequence and the corresponding position in the 3D structure are highlighted in purple. The surface part is colored green, and the transmembrane part is gray.

To experimentally validate the capacity of these regions to be targeted by antibodies, we synthesized peptides covering them. We screened serum samples of lymphoma patients for IgG antibodies binding to them. Of the five regions, we could synthesize peptides for four of them (named peptides 2-5). Among 150 serum samples tested by ELISA at a single dilution, all the peptides showed positive hits: peptide 2 in 46/150 cases (30%), peptide 3 in 49/150 (33%), peptide 4 in 26/150 (17%), and peptide 5 in 15/150 (10%) (Figure 6A, B, C). No significant association has been observed with lymphoma histology except mantle cell lymphoma, which shows a meager percentage of positivity. No association with gender or age has been observed (Figure 6BV, C, D; Table S4).

**Figure 6.**
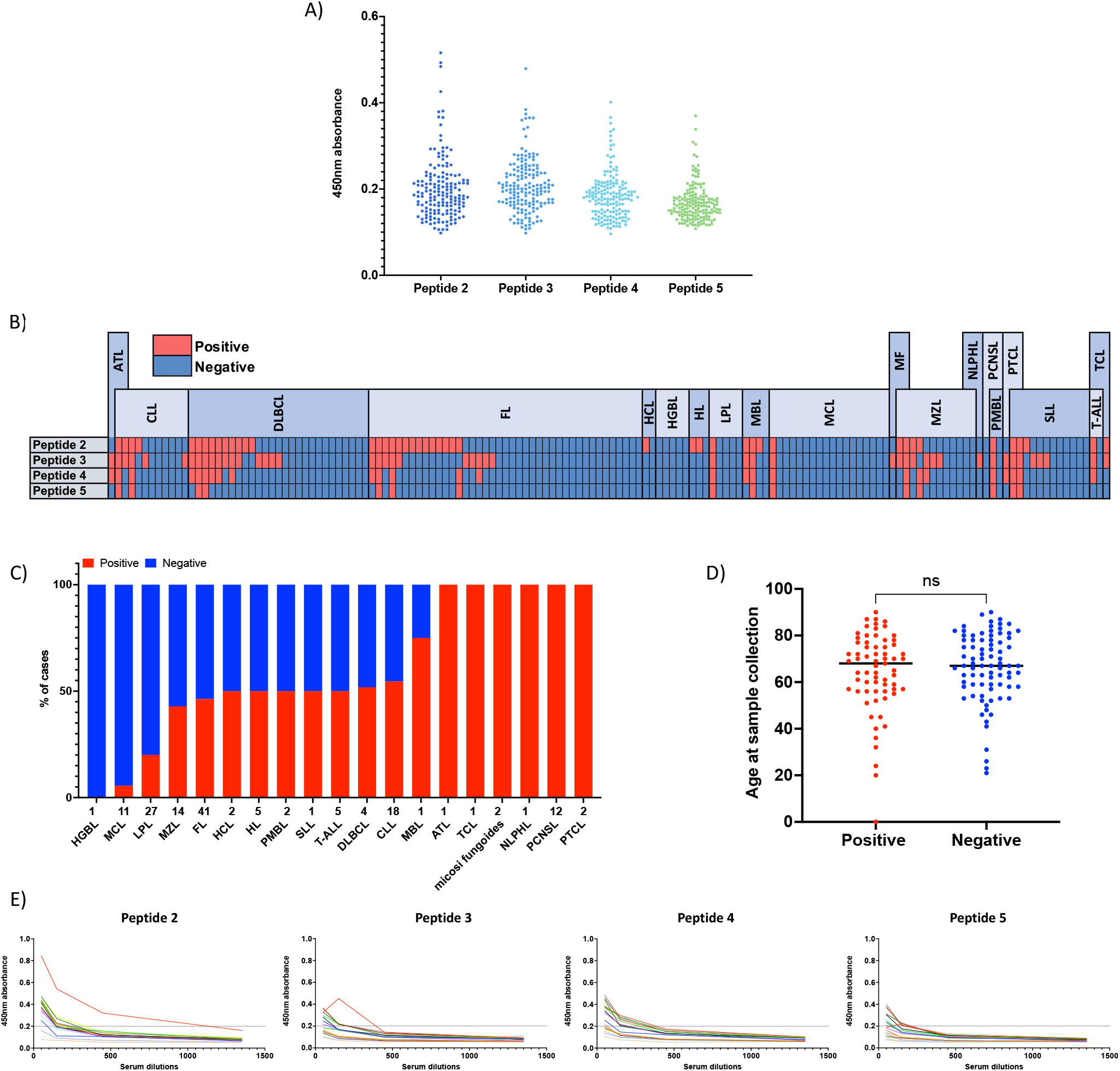
The serum of lymphoma patients contains antibodies targeting the HERV-K envelope. **(A)** ELISA was performed on serum samples from lymphoma patients against the four identified peptides. Y-axes, 450nm absorbance. **(B)** Heatmap showing positive (red) and negative (blue) patients to antibodies against HERV-envelope peptides. **(C)** percentage of positive cases in the different lymphoma subtypes. **(D)** Differences in age at the sample collection between positive and negative patients. **(E)** ELISA validations were performed with serial serum dilutions. ATL, Adult T-cell leukemia; CLL, Chronic lymphocytic leukemia; DLBCL, Diffuse large B-cell lymphoma; FL, Follicular lymphoma; HCL, Hairy cell leukemia; HGBL, High grade B-cell lymphoma; HL, Hodgkin lymphoma; LPL, Lymphoplasmacytic lymphoma; MBL, Monoclonal B-cell lymphocytosis; MCL, Mantel cell lymphoma; MF, mycosis fungoides; MZL, Marginal zone lymphoma; NLPHL, Nodular lymphocyte-predominant Hodgkin lymphoma; PCNSL, Primary CNS Lymphoma; PMBL, Primary mediastinal B-cell lymphoma; PTCL, Peripheral T-Cell Lymphoma; SLL, Small lymphocytic leukemia; T-ALL, T-cell acute lymphoblastic leukemia; TCL, T-cell lymphomas

The binding data were confirmed by serum serial dilution of the most reactive samples in the screening (Figure 6E).

These data indicated that the HERV-K env can be immunogenic and that there are anti-HERV-K env circulating antibodies in the sera of lymphoma patients.

### Novel humanized camelid anti-HERV-K env nanobodies

After proving the immunogenicity of the four peptides, we used them in a phage display screening to identify potential camelid nanobodies capable of recognizing the HERV-K env protein.

We identified two nanobodies that we termed FF-00 and FF-01, both showing binding specificity for peptide 3. The binding affinities of the nanobodies identified during the screening process were validated using surface plasmon resonance (SPR) experiments. Only one strongly interacted with peptide 3 (FF-01) (Figure 7A). This nanobody demonstrated a remarkable binding affinity, with a dissociation constant (Kd) of 142 nM. In contrast, the other nanobody (FF-00) displayed weak or negligible affinity towards the target peptides. FF-00 and FF-01 were humanized with the expression of IgG1 human Fc.

**Figure 7.**
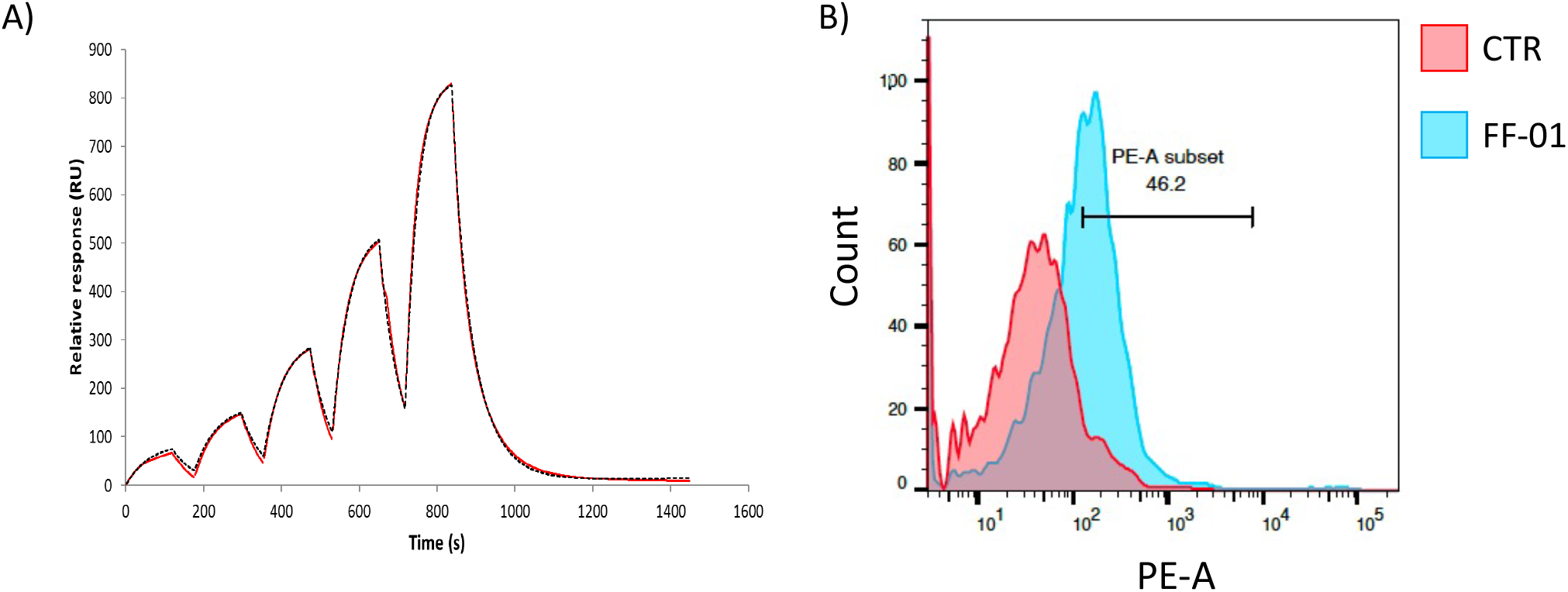
FF-01 binds HERV-K envelope. (A) FF-01 binding data was obtained by surface plasmon resonance on immobilized peptide 3 of the HERV-K envelope. (B) FACS data of FF-01 binding to the extracellular envelope in RCK-8 cell line.

According to SPR data and similarly to the rabbit commercial antibody, FF-01 could bind the env protein on the cellular surface(Figure 7B). At the same time, FF-00 showed only a mild or null binding (data not indicated). The amino acid sequence recognized by FF-01 (peptide 3) was shown to be conserved among different HERV-K family members ^43^. This makes the antibody capable of binding to env proteins coded by other members of the HERV-K family.

### The humanized camelid anti-HERV-K env nanobody FF-01 has anti-lymphoma activity

We first tested the possible direct anti-lymphoma activity of the two nanobodies, FF-00 and FF-01. Still, we did not see any activity superior to the negative control (buffer only) (Figure S6). We then explored the antibody-dependent cellular cytotoxicity (ADCC) potential of FF-00 and FF-01. FF-01 induced ADCC at one μg/ml in the high env expression cell line and not in the remaining cell lines. No sign of ADCC induction was observed with FF-00, and a second negative control antibody targeting protein from Sars-COVID was not expressed in our cell lines (Figure 8A).

**Figure 8.**
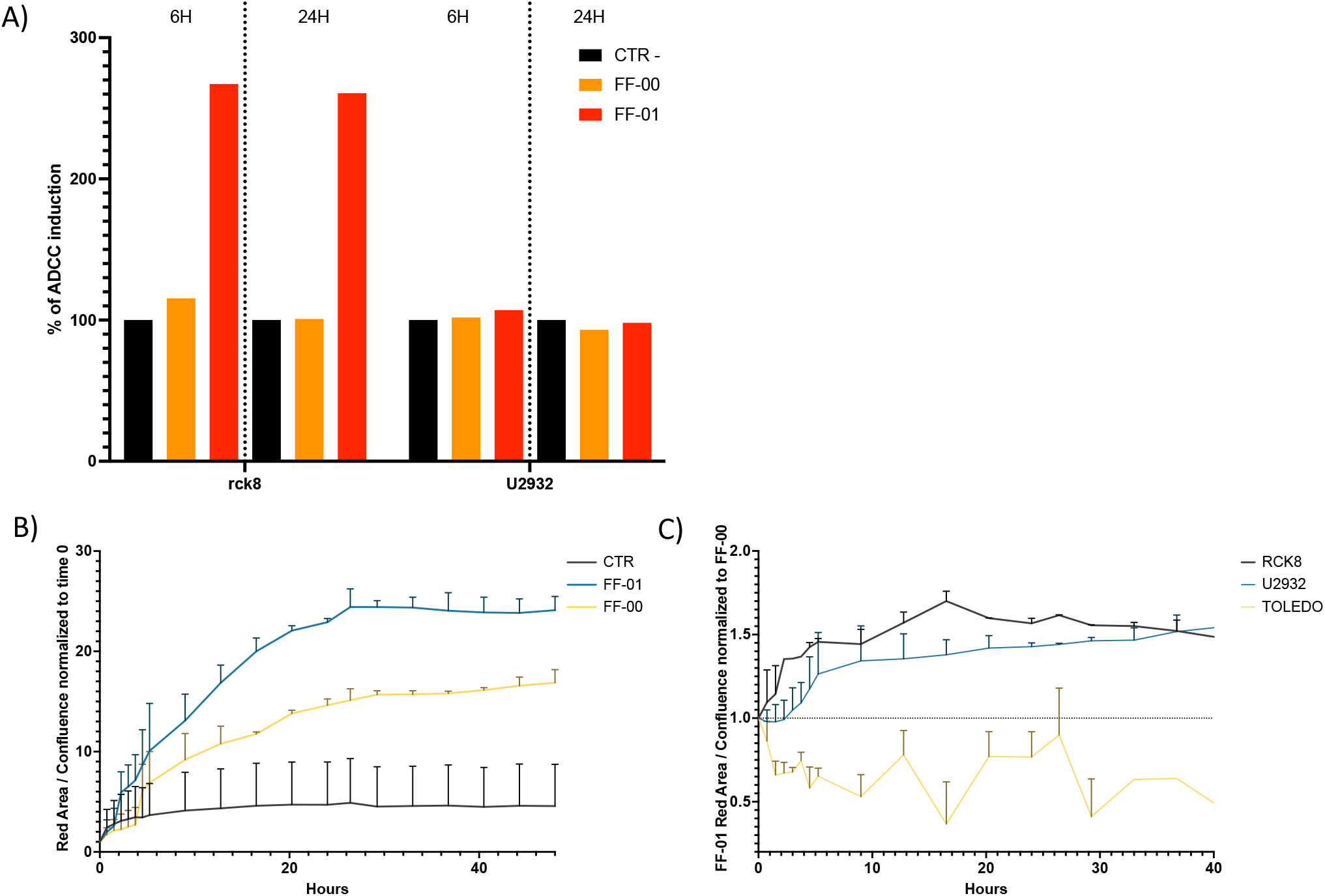
Antibody-dependent cell-mediated Cytotoxicity experiment. **(A)** ADCC performed co-culturing lymphoma cell line RCK8 or U2932 with effector cells in the presence of negative IgG control (CTR-), non-binder Ab (FF-00), and envelope binder antibody (FF-01) for 6 and 24 hours. (B) FF-01 internalization in RCK-8 cell lines. **(B)** FF-01 internalization in RCK-8, U2932, and Toledo cell lines normalized on FF-00 internalization. Cell lines incubated with FF-01 or neg CTR and FabFluor were followed for 40 hours with live cell imaging Incucyte. The increase in red fluorescent area was monitored over time and normalized on confluence.

Finally, as proof of principle for the possible use of the antibody as part of an antibody-drug conjugate (ADC), we analyzed the internalization properties of FF-01 and FF-00. In line with the two antibodies’ binding strength, FF-01 was rapidly internalized in the RCK8, a cell line with high env expression, and, to a lesser extent, in the U2932 expressing low env levels. No internalization was observed in the env-negative TOLEDO. FF-00 was internalized only in the cell line with high env expression (RCK8) but with a much lower signal than FF-01 (Figure 8B).

These data demonstrate the possibility of targeting endogenous retroviruses with antibodies in HERK-env expression cell lines.

## Discussion

We have demonstrated the presence and expression of endogenous retroviruses in lymphoma patients and cell lines. With an ad-hoc analysis, we have identified the expression of transposable elements comprising human endogenous retroviruses in lymphoma patients using reads that are usually discarded from standard RNA-seq data. We separated them into four clusters with different prognoses and recapitulated ABC vs GCB differences. Human endogenous retroviruses (HERV) were differentially expressed in lymphoma patients compared to normal samples, and especially from the family, K HML-2 was overexpressed in tumoral samples compared to normal samples. These data are in line with previously published data ^23^.

The overexpression of pathways related to Interferon, Toll-like receptor, interleukins, and JAK/STAT in patients overexpressing HERV-K113 compared to low expression samples suggests a possible activation of the so-called “viral mimicry” state. Activation of viral mimicry has been explored as a potential intrinsic adjuvant that could sensitize cancer cells for immune recognition, boosting immunotherapeutic strategies ^48^

In lymphoma cell lines, we have confirmed the RNA expression of HERVs, specifically the envelope gene, and demonstrated that the RNA expression could also be translated to protein.

In line with evidence that the HERV env can be expressed on the surface of cancer cell lines ^49^, thereby providing a potential source of new tumor-associated antigens, we demonstrated that HERV env proteins were indeed expressed on the cell surface of lymphoma cell lines. Among the HERVs of the family K, K113 is one of the more recent ones with an intact ORF. We identified one cell line, RCK8, with the K113 heterozygous insertion, while the other tested were negative. Despite the presence of K113 only in RCK8, all the other K113 negative cell lines showed HERVs RNA and protein expression at different degrees, suggesting that K113 is not the only player. Recent evidence has demonstrated that HERV overexpression in cancer is not solely the outcome of epigenetic reshaping without genuine involvement in cancer processes; instead, it appears to be directly implicated in tumor pathogenesis and survival ^50^. By silencing the env RNA expression, we have demonstrated that the HERV envelope sustains the growth of lymphoma cells. Cell lines with downregulation of env showed significantly decreased cellular proliferation compared to control-treated cells.

Since surface expression of the env protein was proven in lymphoma cell lines, we identified five extracellular epitopes (1, 2, 3, 4, and 5) via computational methods. We were able to synthesize four of them (2,3,4,5). Peptides 2 and 3 partially overlapped with the sequence of the commercial rabbit antibody used previously for FACS analysis, confirming their capabilities to be recognized by antibodies.

We confirmed the immunogenicity of the peptides and identified lymphoma patients expressing antibodies able to recognize endogenous retroviruses, thanks to ELISA screening performed in almost 160 lymphoma patients. These patients could be a source of fully humanized antibodies. The presence of anti-HERV antibodies has also been reported in serum samples of patients with Systemic Lupus Erythematosus and breast cancer ^51,52^

We then performed a phage display screening, identifying two camelid nanobodies that recognize peptide 3. The two nanobodies were called FF-00 and FF-01. Surface plasmon resonance experiments confirmed the binding of FF-01 specifically to peptide three and weak/negative binding for FF-00, which was then used as a negative control in the experiments. Nanobodies were coupled with human Fc IgG1, producing a heavy chain antibody. One of the critical advantages of heavy chain antibodies is their smaller size, allowing them to penetrate tissues and tumors more rapidly and intensely than mAbs ^53^. FF-01 also binds to the env complete protein when extracellularly expressed by cell lines.

Recently, antibodies targeting endogenous retroviruses have been shown to promote lung cancer immunotherapy ^49^. To target HERVs, we tested FF-00 and FF-01 for the capability of having a direct anti-lymphoma activity. Both antibodies did not show any direct antitumoral activity. On the contrary, FF-01 can induce antibody-dependent cellular cytotoxicity in line with previous findings in lung cancer ^49^. FF-01 and, to a lesser extent, FF-00 demonstrated internalization in cell lines expressing the env protein on their surface.

These data demonstrate the expression of human endogenous retroviruses in DLBCL patients and propose them as a new therapy target for lymphomas.

## Author’s contribution

F. Spriano designed the study, coordinated the project, performed experiments, analyzed the data, and wrote the paper. F. Bertoni designed the study, coordinated the project, analyzed the data, provided resources, and wrote the paper. L. Cascione performed bioinformatic analysis and wrote the paper. J. Sgrignani performed experiments and analyzed the data. N. Bendik, S. Napoli, G. Sartori, E. Cannas, T. Gong, A. J. Arribas, and M. Pizzi performed experiments. D. Rossi provided well-characterized clinical specimens, and provided advice. D. F. Robbiani and A. Cavalli provided resources and advice. All authors read and approved the final manuscript.

