## Supplementary tables and figures for "Characterization of a novel humanized heavy chain antibody targeting endogenous retroviruses with anti-lymphoma activity"

### Supplementary figures and tables

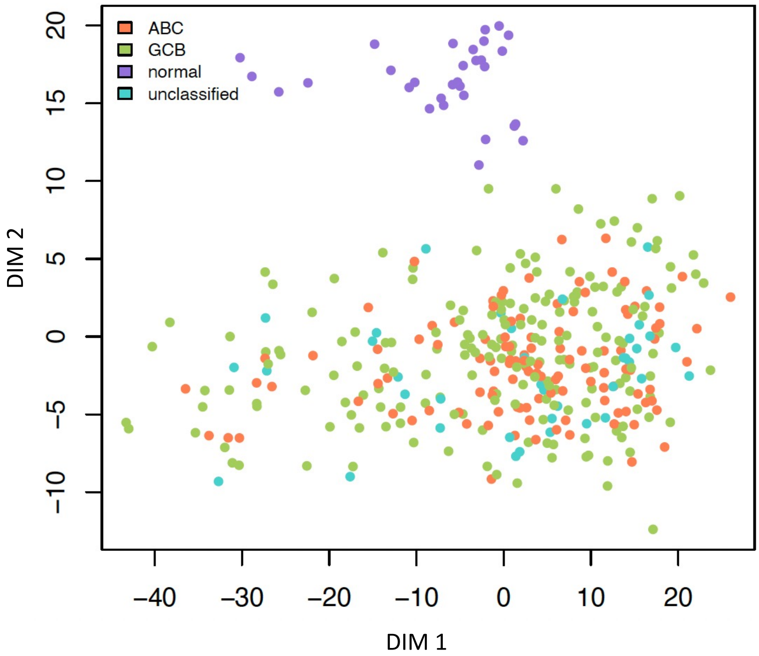


**Supplementary Figure 1. Multidimensional scaling (MDS) plot of HML-2 HERVs on lymphoma patients**. Multidimensional scaling (MDS) plot showing normal and tumoral samples clustering differently based on HML-2 expression.

| | NAME | SIZE | NES | NOM p-val | FDR q-val |
|---|---|---|---|---|---|
| Antigen Presentation and Processing | REACTOME_ANTIGEN_PRESENTATION_FOLDING_ASSEMBLY_AND_PEPTIDE_LOADING_OF_CLASS_I_MHC | 28 | 2.00 | 0.000 | 0.002 |
| | REACTOME_CLASS_I_MHC_MEDIATED_ANTIGEN_PROCESSING_PRESENTATION | 341 | 1.93 | 0.000 | 0.004 |
| | REACTOME_ANTIGEN_PROCESSING_UBIQUITINATION_PROTEASOME_DEGRADATION | 277 | 1.84 | 0.000 | 0.009 |
| Viral Infection | WP_HEPATITIS_B_INFECTION | 130 | 2.59 | 0.000 | 0.000 |
| | WP_MEASLES_VIRUS_INFECTION | 112 | 2.36 | 0.000 | 0.000 |
| | WP_TCELL_ANTIGEN_RECEPTOR_TCR_PATHWAY_DURING_STAPHYLOCOCCUS_AUREUS_INFECTION | 60 | 2.18 | 0.000 | 0.000 |
| | REACTOME_EXPORT_OF_VIRAL_RIBONUCLEOPROTEINS_FROM_NUCLEUS | 29 | 2.18 | 0.000 | 0.000 |
| | REACTOME_LATE_SARS_COV_2_INFECTION_EVENTS | 64 | 1.99 | 0.000 | 0.002 |
| | KEGG_LEISHMANIA_INFECTION | 69 | 1.94 | 0.000 | 0.003 |
| | WP_ACUTE_VIRAL_MYOCARDITIS | 82 | 1.92 | 0.000 | 0.004 |
| | WP_SARS_CORONAVIRUS_AND_INNATE_IMMUNITY | 14 | 1.91 | 0.000 | 0.004 |
| | REACTOME_SARS_COV_1_HOST_INTERACTIONS | 88 | -0.57 | 0.000 | 0.000 |
| | REACTOME_INFLUENZA_INFECTION | 147 | -0.65 | 0.000 | 0.000 |
| | REACTOME_SARS_COV_2_MODULATES_HOST_TRANSLATION_MACHINERY | 49 | -0.79 | 0.000 | 0.000 |
| | REACTOME_SARS_COV_1_MODULATES_HOST_TRANSLATION_MACHINERY | 35 | -0.87 | 0.000 | 0.000 |
| | REACTOME_SARS_COV_1_INFECTION | 131 | -1.98 | 0.000 | 0.006 |
| Viral Infection / Interferon | WP_TYPE_I_INTERFERON_INDUCTION_AND_SIGNALING_DURING_SARSCOV2_INFECTION | 25 | 2.16 | 0.000 | 0.000 |
| | REACTOME_ANTIVIRAL_MECHANISM_BY_IFN_STIMULATED_GENES | 76 | 2.03 | 0.000 | 0.001 |
| Interferon | WP_HOSTPATHOGEN_INTERACTION_OF_HUMAN_CORONAVIRUSES_INTERFERON_INDUCTION | 30 | 2.21 | 0.000 | 0.000 |
| | REACTOME_INTERFERON_SIGNALING | 171 | 2.21 | 0.000 | 0.000 |
| | WP_INTERFERON_TYPE_I_SIGNALING_PATHWAYS | 52 | 2.18 | 0.000 | 0.000 |
| | REACTOME_INTERFERON_GAMMA_SIGNALING | 81 | 2.16 | 0.000 | 0.000 |
| | HALLMARK_INTERFERON_ALPHA_RESPONSE | 94 | 2.14 | 0.000 | 0.000 |
| | HALLMARK_INTERFERON_GAMMA_RESPONSE | 192 | 2.11 | 0.000 | 0.000 |
| | REACTOME_INTERFERON_ALPHA_BETA_SIGNALING | 58 | 2.10 | 0.000 | 0.001 |
| | WP_TYPE_II_INTERFERON_SIGNALING | 32 | 2.04 | 0.000 | 0.001 |
| | WP_OVERVIEW_OF_INTERFERONSMEDIATED_SIGNALING_PATHWAY | 16 | 1.92 | 0.004 | 0.004 |
| | REACTOME_REGULATION_OF_IFNA_IFNB_SIGNALING | 12 | 1.85 | 0.004 | 0.008 |
| JAK/STAT | KEGG_JAK_STAT_SIGNALING_PATHWAY | 107 | 2.01 | 0.000 | 0.002 |
| | WP_REGULATORY_CIRCUITS_OF_THE_STAT3_SIGNALING_PATHWAY | 70 | 1.96 | 0.000 | 0.003 |
| Interleukin | WP_IL1_SIGNALING_PATHWAY | 52 | 2.34 | 0.000 | 0.000 |
| | PID_IL2_1PATHWAY | 55 | 2.33 | 0.000 | 0.000 |
| | WP_INTERLEUKIN1_IL1_STRUCTURAL_PATHWAY | 49 | 2.28 | 0.000 | 0.000 |
| | SIG_IL4RECEPTOR_IN_B_LYPHOCYTES | 27 | 2.27 | 0.000 | 0.000 |
| | WP_IL4_SIGNALING_PATHWAY | 53 | 2.26 | 0.000 | 0.000 |
| | WP_IL3_SIGNALING_PATHWAY | 45 | 2.24 | 0.000 | 0.000 |
| | REACTOME_INTERLEUKIN_2_FAMILY_SIGNALING | 37 | 2.24 | 0.000 | 0.000 |
| | BIOCARTA_IL6_PATHWAY | 21 | 2.17 | 0.000 | 0.000 |
| | WP_IL2_SIGNALING_PATHWAY | 41 | 2.13 | 0.000 | 0.000 |
| | PID_IL2_STAT5_PATHWAY | 30 | 2.11 | 0.000 | 0.000 |
| | BIOCARTA_IL2_PATHWAY | 22 | 2.09 | 0.000 | 0.001 |
| | BIOCARTA_IL2RB_PATHWAY | 37 | 2.06 | 0.000 | 0.001 |
| | WP_IL9_SIGNALING_PATHWAY | 14 | 2.06 | 0.000 | 0.001 |
| | REACTOME_INTERLEUKIN_17_SIGNALING | 66 | 2.05 | 0.000 | 0.001 |
| | WP_INTERLEUKIN11_SIGNALING_PATHWAY | 41 | 2.04 | 0.000 | 0.001 |
| | REACTOME_INTERLEUKIN_15_SIGNALING | 14 | 2.03 | 0.000 | 0.001 |
| | REACTOME_INTERLEUKIN_3_INTERLEUKIN_5_AND_GM_CSF_SIGNALING | 43 | 2.01 | 0.000 | 0.001 |
| | BIOCARTA_IL3_PATHWAY | 13 | 2.00 | 0.000 | 0.002 |
| | PID_IL1_PATHWAY | 33 | 1.99 | 0.000 | 0.002 |
| | REACTOME_INTERLEUKIN_RECEPTOR_SHC_SIGNALING | 22 | 1.98 | 0.000 | 0.002 |
| | WP_IL17_SIGNALING_PATHWAY | 25 | 1.98 | 0.000 | 0.002 |
| | WP_IL7_SIGNALING_PATHWAY | 24 | 1.98 | 0.002 | 0.002 |
| | BIOCARTA_IL1R_PATHWAY | 29 | 1.94 | 0.000 | 0.003 |
| | REACTOME_FCGR3A_MEDIATED_IL10_SYNTHESIS | 86 | 1.94 | 0.000 | 0.003 |
| | WP_IL5_SIGNALING_PATHWAY | 38 | 1.91 | 0.000 | 0.004 |
| | WP_IL6_SIGNALING_PATHWAY | 39 | 1.91 | 0.002 | 0.004 |
| | PID_IL12_2PATHWAY | 59 | 1.91 | 0.000 | 0.004 |
| | REACTOME_INTERLEUKIN_20_FAMILY_SIGNALING | 18 | 1.90 | 0.004 | 0.005 |
| | BIOCARTA_IL7_PATHWAY | 16 | 1.82 | 0.003 | 0.010 |
| | REACTOME_INTERLEUKIN_6_SIGNALING | 11 | 1.82 | 0.003 | 0.010 |
| Toll Like Receptor | WP_TOLLLIKE_RECEPTOR_SIGNALING_PATHWAY | 82 | 2.25 | 0.000 | 0.000 |
| | KEGG_TOLL_LIKE_RECEPTOR_SIGNALING_PATHWAY | 81 | 2.21 | 0.000 | 0.000 |
| | BIOCARTA_TOLL_PATHWAY | 23 | 2.16 | 0.000 | 0.000 |
| | REACTOME_TOLL_LIKE_RECEPTOR_CASCADES | 155 | 2.13 | 0.000 | 0.000 |
| | WP_TOLLLIKE_RECEPTOR_SIGNALING_RELATED_TO_MYD88 | 28 | 2.13 | 0.000 | 0.000 |
| | KEGG_FC_GAMMA_R_MEDIATED_PHAGOCYTOSIS | 85 | 2.12 | 0.000 | 0.000 |
| | REACTOME_MYD88_INDEPENDENT_TLR4_CASCADE | 106 | 2.11 | 0.000 | 0.001 |
| | REACTOME_TOLL_LIKE_RECEPTOR_9_TLR9_CASCADE | 104 | 2.10 | 0.000 | 0.001 |
| | REACTOME_TOLL_LIKE_RECEPTOR_TLR1_TLR2_CASCADE | 108 | 2.08 | 0.000 | 0.001 |
| | WP_TLR4_SIGNALING_AND_TOLERANCE | 25 | 2.02 | 0.000 | 0.001 |
| B_T cell receptor | WP_B_CELL_RECEPTOR_SIGNALING_PATHWAY | 92 | 2.55 | 0.000 | 0.000 |
| | WP_TCELL_RECEPTOR_SIGNALING_PATHWAY | 84 | 2.53 | 0.000 | 0.000 |
| | KEGG_T_CELL_RECEPTOR_SIGNALING_PATHWAY | 101 | 2.50 | 0.000 | 0.000 |
| | PID_CD8_TCR_PATHWAY | 49 | 2.47 | 0.000 | 0.000 |
| | PID_TCR_PATHWAY | 59 | 2.45 | 0.000 | 0.000 |
| | SIG_BCR_SIGNALING_PATHWAY | 45 | 2.36 | 0.000 | 0.000 |
| | KEGG_B_CELL_RECEPTOR_SIGNALING_PATHWAY | 71 | 2.28 | 0.000 | 0.000 |
| | WP_T_CELL_RECEPTOR_AND_COSTIMULATORY_SIGNALING | 28 | 2.07 | 0.002 | 0.001 |
| | PID_CD8_TCR_DOWNSTREAM_PATHWAY | 53 | 2.06 | 0.000 | 0.001 |
| | BIOCARTA_BCR_PATHWAY | 33 | 1.83 | 0.002 | 0.009 |
| | REACTOME_ANTIGEN_ACTIVATES_B_CELL_RECEPTOR_BCR_LEADING_TO_GENERATION_OF_SECOND_MESSENGERS | 79 | 2.09 | 0.000 | 0.001 |

**Supplementary Figure 2. Gene-sets enrichment analysis comparing K113 high v low expression patients.** Positive NES identifies gene sets overexpressed in high K113 expression NES = normalized enrichment score.

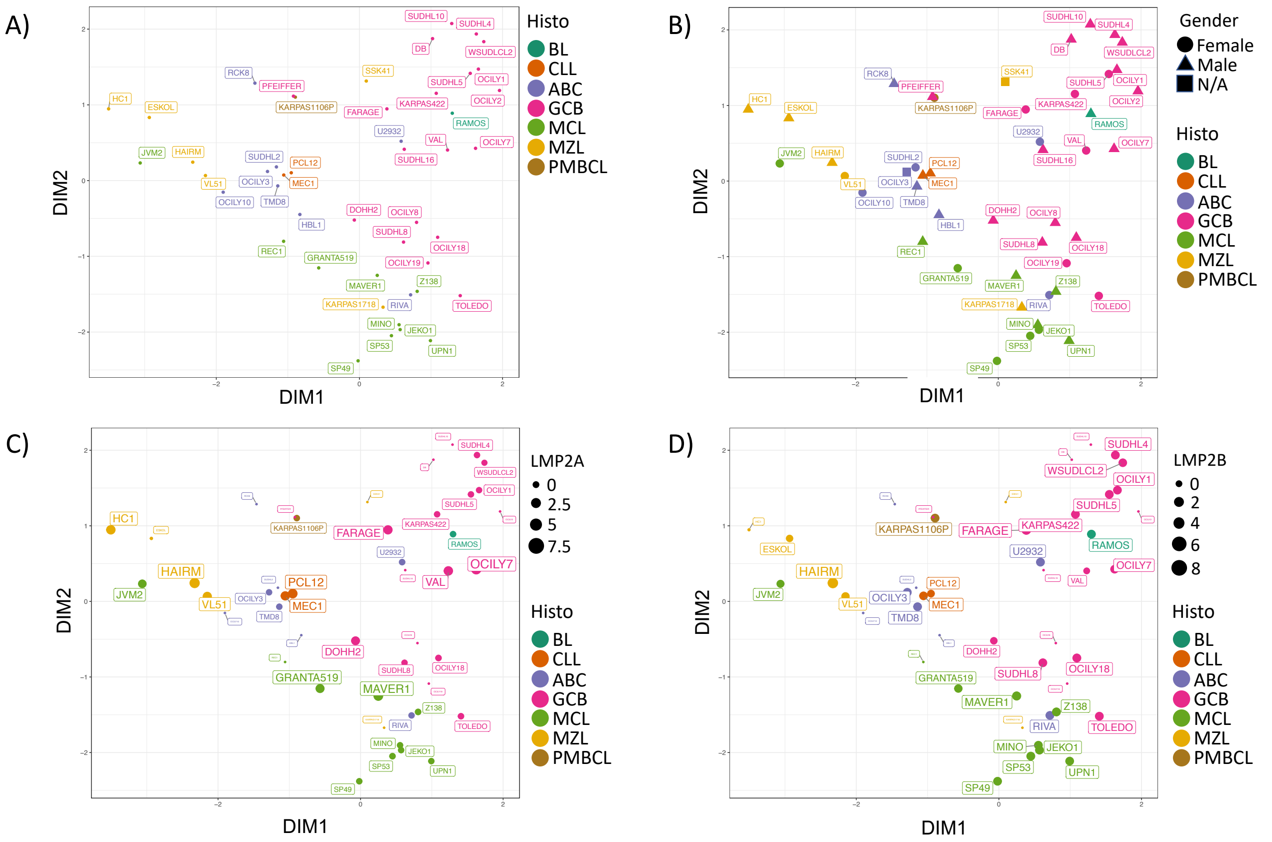


**Supplementary Figure 3. Multidimensional scaling (MDS) plot on cell lines**. (A) MDS plot on cell lines divided by subtype. (B) MDS plot on cell lines divided by subtype and gender. (C-D) MDS plot on cell lines divided by subtype and EBV expression (LMP2A and LMP2B).

RCK8

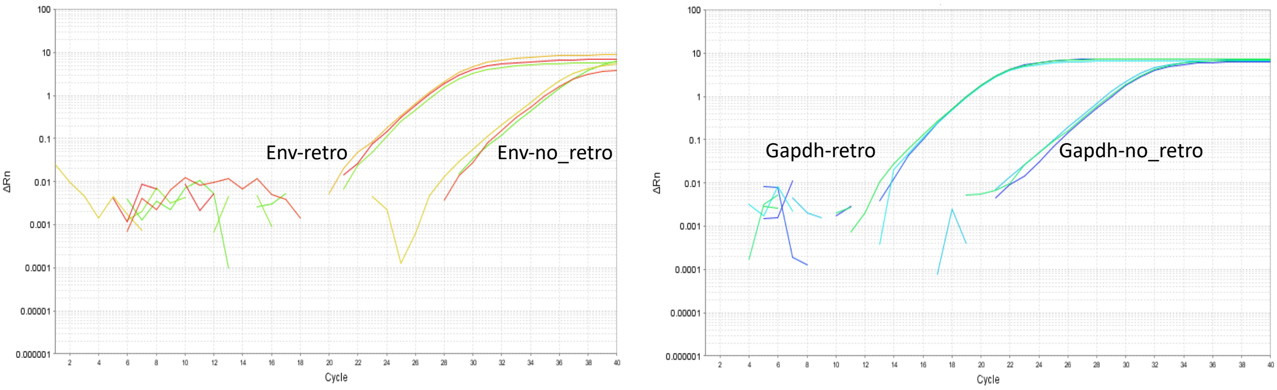


**Supplementary Figure 4. RNA envelope expression in RCK8 cell line.** Representative graph of HERV-K envelope expression by qPCR in RCK8 cell line. Retrotranscribed (retro) samples were compared to non-retrotranscribed (no-retro) samples.

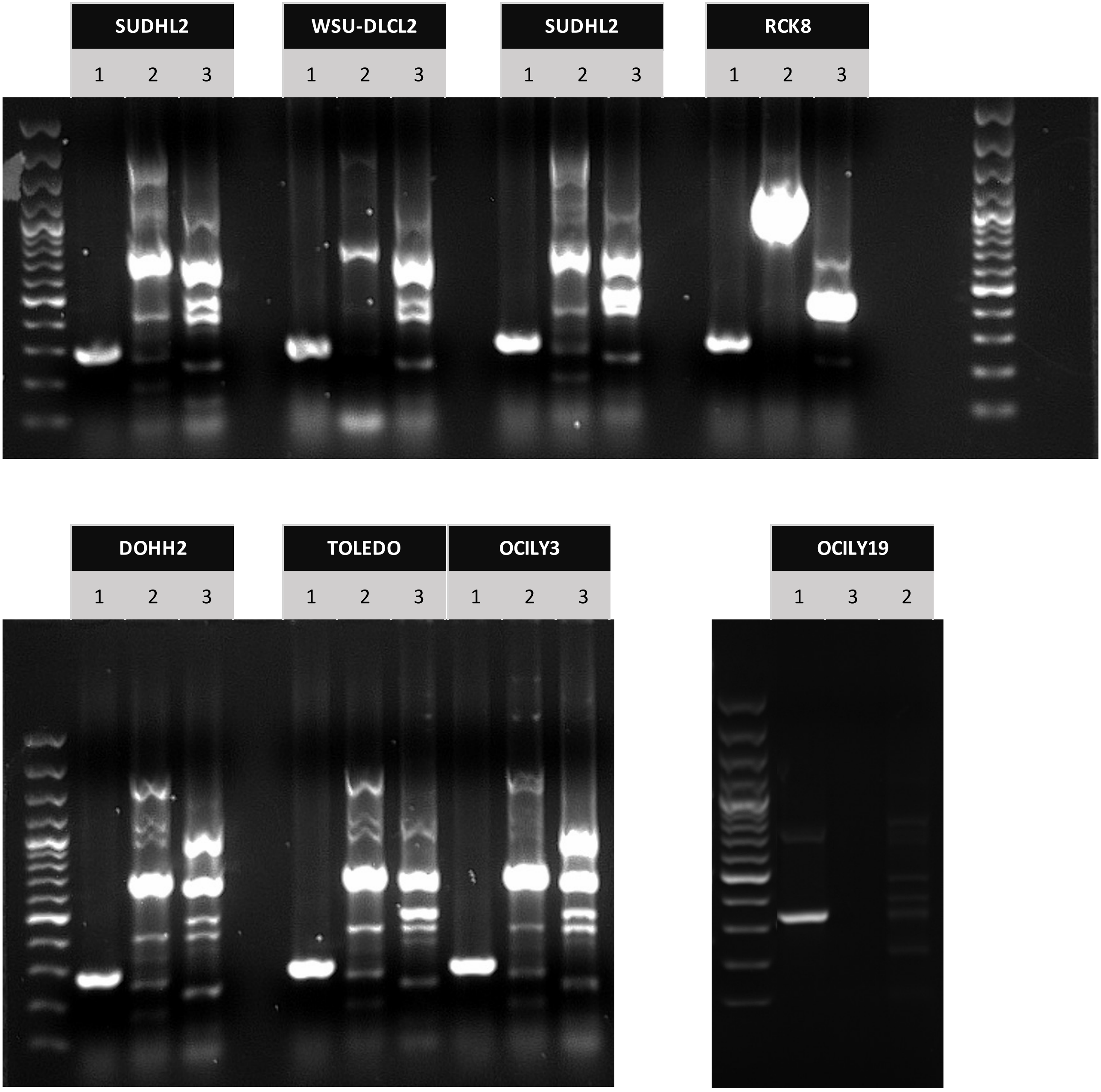


**Supplementary Figure 5. The pattern of K113 insertion in the genome.** Agarose gel electrophoresis after PCR showed a full-length heterozygous insertion of HERV-K113 in RCK8. Lane 1, preintegration site; lane 2, K113 5' LTR; lane 3, K113 3' LTR.

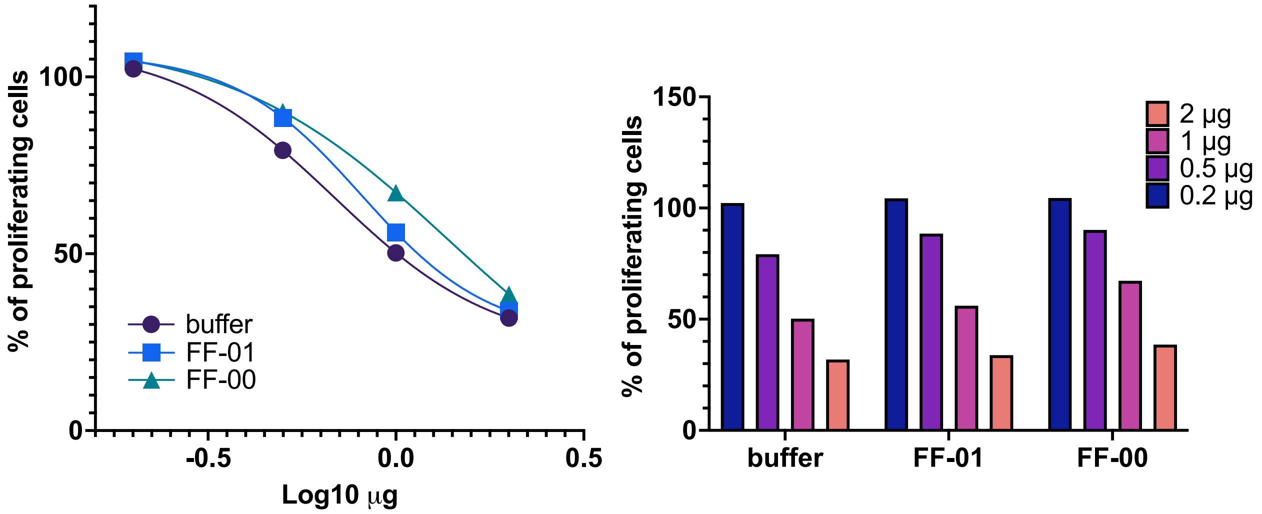


**Supplementary Figure 6. Anti-proliferation effect of FF-01**. RCK8 cell lines were treated for 72 hours with increasing anti-HERV-K envelope antibody FF-01.

| | Cluster_1 | Cluster_2 | Cluster_3 | Cluster_4 |
|---|---|---|---|---|
| NEG | 58 | 29 | 91 | 88 |
| POS | 26 | 13 | 15 | 13 |
| UNCLASS | 2 | 0 | 1 | 1 |

**Supplementary Table 1. Patient distribution in the four clusters is based on the double hit/dark zone signature.**

| | **sense siRNA (5p --> 3p)** | **antisense siRNA (5p --> 3p)** | **Starting target position** | **Ending target position** |
|---|---|---|---|---|
| **siRNA 2** | GCAGCUAACUAUACCUACUTT | AGUAGGUAUAGUUAGCUGCTT | 232- | 250 |
| **siRNA 3** | CGAGGUCAAUUCUACCACATT | UGUGGUAGAAUUGACCUCGTT | 742- | 760 |

**Supplementary Table 3. siRNA sequences and target binding position.**
